# NREM sleep microarchitecture is associated with internalizing symptoms and emotional functioning in early adolescence

**DOI:** 10.64898/2026.10.04.756568

**Authors:** Hazal Arpaci, Morgan Lott, Sarah Dickens, Aditi Tripathy, Nandita Banik, Intisar Becic, Narges Firouzshahi, Anna Holowicki, Candice Alfano, Gerta Bardoshi, Bengi Baran

## Abstract

**Study Objectives:** Sleep disturbances are common in youth with anxiety disorders, yet most prior work relies on subjective measures and does not capture underlying neural processes. We examined whether alterations in non-rapid eye movement (NREM) sleep oscillations are present in children with anxiety disorders and whether these features relate to internalizing symptoms and emotional functioning.

**Methods:** Children aged 9-13 years with diagnosed anxiety disorders (ANX; n = 17) and typically developing non-psychiatric comparison participants (NC; n = 26) completed overnight polysomnography. Slow oscillations (SO) and slow and fast sleep spindles were characterized using validated automated detection algorithms. Participants also completed a pre-sleep emotional processing task, and parents provided standardized measures of youth mental health. Group differences and dimensional associations between NREM oscillations, internalizing symptoms, and behavioral indices of emotional processing were examined.

**Results:** Sleep architecture did not differ between groups. Slow spindle duration was shorter in the ANX group. Across the full sample, shorter slow spindle duration was associated with greater internalizing symptoms, whereas higher SO density was positively associated with internalizing symptoms. Children in the ANX group exhibited greater facial emotional arousal and more frequent look-away behavior during the emotional processing task. Across participants, greater emotional arousal and look-away behavior were associated with shorter spindle duration during subsequent sleep.

**Conclusions:** Specific features of NREM microarchitecture were associated with individual differences in internalizing symptoms and emotional functioning during early adolescence. These findings highlight the potential relevance of NREM oscillations for understanding developmental links between sleep and youth psychopathology.

**Statement of Significance:** Sleep disturbances are common in youth with anxiety disorders, yet most research has relied on subjective reports and broad measures of sleep quality. This study demonstrates that specific features of non-rapid eye movement sleep microarchitecture, particularly sleep spindle duration, are associated with anxiety, depression, and emotional responding during early adolescence even in the absence of differences in overall sleep quality or sleep stage distribution. These findings highlight the importance of examining neural features of sleep when investigating emotional functioning and internalizing psychopathology during development. More broadly, the results suggest that adolescence may represent a sensitive developmental window during which alterations in sleep physiology are linked to emotional functioning. Longitudinal studies are needed to determine how these sleep features relate to developmental trajectories of youth psychopathology.

## Introduction

Sleep plays a vital role in emotional functioning and mental health. Across development, adequate and efficient sleep contributes to the processing and regulation of emotional experiences^1–3^, while disturbances in sleep are associated with heightened emotional reactivity^4,5^ and vulnerability to internalizing symptoms^6,7^. The transition from late childhood to early adolescence is a period of pronounced changes in sleep timing and duration^8,9^ and the morphology of non-rapid eye movement (NREM) sleep oscillations. This developmental period is also associated with an increased likelihood of the emergence of anxiety disorders^10^, providing a valuable opportunity to examine whether alterations in sleep physiology are detectable when anxiety becomes clinically identifiable. Despite these overlapping developmental processes, relatively little is known about how NREM oscillatory activity relates to emotional functioning and anxiety during early adolescence. The present study examines whether NREM sleep oscillations are altered in youth with anxiety disorders and whether these alterations are associated with internalizing symptoms and behavioral indices of emotional processing.

Generalized Anxiety Disorder (GAD), Separation Anxiety Disorder (SAD), and Social Anxiety Disorder (social phobia; SOC) are among the most commonly observed anxiety disorders in adolescence and, due to substantial overlap in clinical presentation and pathophysiology, have been collectively referred to as the “pediatric anxiety triad”^11^. These disorders often persist into adulthood^12–15^. Further, depressive symptoms commonly co-occur with pediatric anxiety disorders and contribute substantially to clinical impairment, justifying the examination of internalizing symptoms more broadly in this population^16,17^.

Anxiety and sleep problems frequently co-occur, potentially reflecting overlapping neurobiological mechanisms. Experimental studies have shown that sleep deprivation increases anxiety the following day^18,19^. Neuromodulatory systems involved in sleep-wake regulation, including adenosinergic signaling, also contribute to anxiety-related processes^20–22^. Adolescence is characterized by substantial maturation of fronto-limbic circuits involved in emotion regulation alongside developmental changes in sleep physiology, raising the possibility that alterations in these processes may be particularly relevant to the emergence of anxiety during this period^23^. In a large pediatric sample, 85% of children and adolescents with anxiety disorders were reported by parents to have concurrent sleep problems^24^. Examples of sleep problems that are faced by this population include delayed bedtime, diminished total sleep time, night waking and nightmares^25–27^. Moreover, sleep disturbances in childhood may predict later anxiety disorders, depressive symptoms and behavioral problems^28–31^. However, most of this literature relies on self-, parent-or clinician-reported sleep duration, sleep quality, or daytime sleepiness. As a result, relatively little is known about whether specific neurophysiological features of sleep are altered in youth with anxiety disorders or how these neural sleep processes relate to emotional functioning and internalizing psychopathology.

Oscillations characteristic of stages 2 and 3 NREM sleep, such as sleep spindles and slow oscillations (SO), reflect coordinated thalamocortical and cortical network activity. A substantial body of work has linked the expression and temporal coordination of these oscillations to emotional memory consolidation^32–37^, suggesting a role for NREM sleep in the integration of emotionally salient experiences. Consistent with this, studies in young adults have shown that greater SO density is associated with reduced emotional reactivity to negative stimuli^38^ and higher slow wave activity (SWA; spectral power in 0.5-4 Hz band) is linked to lower anxiety levels the following morning^39^. In addition, greater spindle activity has been associated with the use of more adaptive emotion regulation strategies in both preschool^40^ and school-aged children^41^. Together, these findings support a broader role for NREM oscillatory activity in affective functioning. In line with this literature, alterations in NREM oscillations have been reported across several psychiatric conditions including schizophrenia^42–44^, bipolar disorder^45^, PTSD^46–48^, anxiety^38^ and depression^49–51^. These findings suggest that NREM sleep microarchitecture may be particularly relevant for understanding sleep-related mechanisms linking sleep and emotional functioning.

Emerging work suggests that NREM oscillatory activity may be altered in youth with anxiety disorders. To date, two studies have reported reduced sleep spindle activity in children and adolescents with anxiety relative to typically developing peers^52,53^. Meers and colleagues did not observe alterations in NREM spindle activity in children with generalized anxiety disorder^52^, whereas Wilhelm and colleagues reported reduced Sigma activity associated with heightened emotional reactivity and greater anxiety symptoms in children and adolescents with social anxiety disorder^53^. Together, these findings raise the possibility that disruptions in NREM oscillations may reflect alterations in neurophysiological processes relevant to emotional regulation and internalizing symptoms during adolescence.

The current study provides an initial test of this possibility by characterizing NREM sleep physiology during a critical developmental window spanning late childhood to early adolescence (9-13 years of age), when anxiety disorders become increasingly prevalent and clinically identifiable. Using overnight polysomnography (PSG) in children with diagnosed anxiety disorders and a typically developing comparison group, we examined whether differences in sleep spindle and SO activity are evident during this developmental stage. In addition, participants completed an emotional processing task prior to sleep, and both parents and children completed standardized measures of youth mental health, enabling examination of emotional regulation and internalizing symptoms in relation to NREM sleep physiology. We hypothesized that children with anxiety disorders would exhibit reduced SO and spindle activity relative to the comparison group, and that greater internalizing symptoms would be associated with alterations in NREM oscillations. Finally, we hypothesized that children with anxiety disorders would exhibit altered emotional regulation, and these behavioral indices would be associated with NREM oscillations and internalizing symptoms.

## Materials and Methods

### Participants

Forty-three children and adolescents (9-13 years of age, 17 males, 26 females) participated in the study (see Table 1 for participant characteristics). The Anxiety group (ANX; n= 17) consisted of youth with a medical chart diagnosis of anxiety disorders (Generalized Anxiety Disorder, Social Anxiety Disorder, or Separation Anxiety Disorder) identified through the University of Iowa Institute for Clinical and Translational Science TrinetX resources and recruited via email or letter invitations. Diagnoses were confirmed with the Kiddie Schedule for Affective Disorders and Schizophrenia-Present and Lifetime Version for DSM-5 (K-SADS-PL^54^). Non-psychiatric comparison subjects (NC; n=26) consisted of typically developing youth who were demographically matched to the anxiety group on age, sex, and mean parental education. Comparison group participants were recruited from the community through electronic flyers and were excluded for any personal history of psychiatric disorders. General exclusion criteria were Body Mass Index (BMI) >30, presence of ADHD or other neurological, metabolic, psychotic or intellectual and developmental disorders, diagnosed sleep disorders (non-diagnosed sleep complaints such as occasional insomnia or inconsistent sleep schedules were not exclusion criteria), a history of a significant head injury that resulted in prolonged loss of consciousness or neurological sequelae, or a family history of psychotic disorders or substance/alcohol use disorders. After the procedures were explained, research staff obtained written informed consent from the parent/caregiver and assent from participants. Families received monetary compensation for their participation. All study procedures were approved by the University of Iowa Human Subjects Office.

**Table 1.** Sample demographics.

|  | Anxiety<br>(n=17) |  | Comparison<br>(n=26) |  | *t* | *p* | Cohen's *d* | 95% CI |
| --- | --- | --- | --- | --- | --- | --- | --- | --- |
|  | *m* | *SD* | *m* | *SD* |  |  |  |  |
| Age (yrs) | 11.3 | 1.6 | 11.3 | 1.4 | 0.1 | .96 | 0.02 | -0.6;0.6 |
| Sex<sup>a</sup> | 10 F/ 7 M | | 16 F/ 10 M | | $\chi^2=.03$ | .86 | | |
| Mean Parental Education (yrs) | 16.4 | 3.1 | 17.0 | 1.7 | -0.7 | .52 | 0.3 | -1.0;0.5 |
| Puberty<sup>b</sup> | 2.2 | 1 | 2.2 | 0.8 | 0.1 | .94 | 0.02 | -0.6;0.7 |
| PDSS<sup>c</sup> | 15.5 | 5.4 | 13.0 | 4.2 | 1.6 | .12 | 0.5 | -0.1;1.2 |
| CHAOS<sup>d</sup> | 3.6 | 3.2 | 3.4 | 2.3 | 0.2 | .87 | 0.1 | -0.6;0.7 |
| PANAS-C<sup>e</sup> |  |  |  |  |  |  |  |  |
| *Positive Affect* | 17.2 | 3.8 | 18.0 | 3.2 | -0.7 | .47 | 0.2 | -0.9;0.4 |
| *Negative Affect* | 13.0 | 5.0 | 9.8 | 3.2 | 2.3 | .03* | 0.8 | 0.1;1.4 |
| RCADS-P<sup>f</sup> |  |  |  |  |  |  |  |  |
| *Total Score* | 32.9 | 11.1 | 13.4 | 8.2 | 6.2 | <.001* | 2.0 | 1.2;2.8 |
| *Depression Score* | 6.1 | 2.9 | 2.1 | 1.8 | 5.1 | <.001* | 1.7 | 0.9;2.4 |
| *Total Anxiety Score* | 23.4 | 8.5 | 10.7 | 6.4 | 5.3 | <.001* | 1.7 | 0.9;2.4 |
| CBCL<sup>g</sup> |  |  |  |  |  |  |  |  |
| *Total problems* | 53.9 | 6.6 | 42.8 | 10.8 | 4.0 | <.001* | 1.3 |  |
| *Internalizing Problems* | 61.6 | 7.5 | 47.6 | 10.9 | 4.8 | <.001* | 1.5 | 0.8;2.2 |
| *Externalizing Problems* | 45.4 | 6.4 | 41.8 | 7.6 | 1.6 | .12 | 0.5 | -0.2;1.2 |

<sup>a</sup>Sex assigned at birth. 2 participants identified as non-binary; all other participants identified as cisgender.
<sup>b</sup>Self-report puberty scale<sup>71</sup>, prepubertal=3, early pubertal=4 and 5;
<sup>c</sup>Pediatric Daytime Sleepiness Scale (PDSS<sup>72</sup>) total score;
<sup>d</sup>Confusion Hubbub and Order Scale (CHAOS<sup>70</sup>) total score;
<sup>e</sup>Positive and Negative Affect Schedule for Children (PANAS-C<sup>69</sup>);
<sup>f</sup>Revised Children's Anxiety and Depression Scale (RCADS<sup>66</sup>);
<sup>g</sup>Child Behavior Checklist (CBCL<sup>67</sup>) t-scores;
\**p*<.05

### Study Design

Participants underwent overnight PSG monitoring, completed an emotional processing task, and participated in clinical interviews (Figure 1A). Children and their primary caregivers arrived at the sleep lab approximately 2 hours prior to the participant’s typical bedtime. Following consent and assent procedures, they completed self and parental-report measures. Participants then completed the emotional processing task in a separate testing room (Figure 1B). This was followed by PSG setup for overnight sleep monitoring in a sound-and light-proofed environment designed to resemble a typical bedroom. Clinical interviews were conducted via videoconference on a separate day approximately 2 weeks after participation.

**Figure 1.**
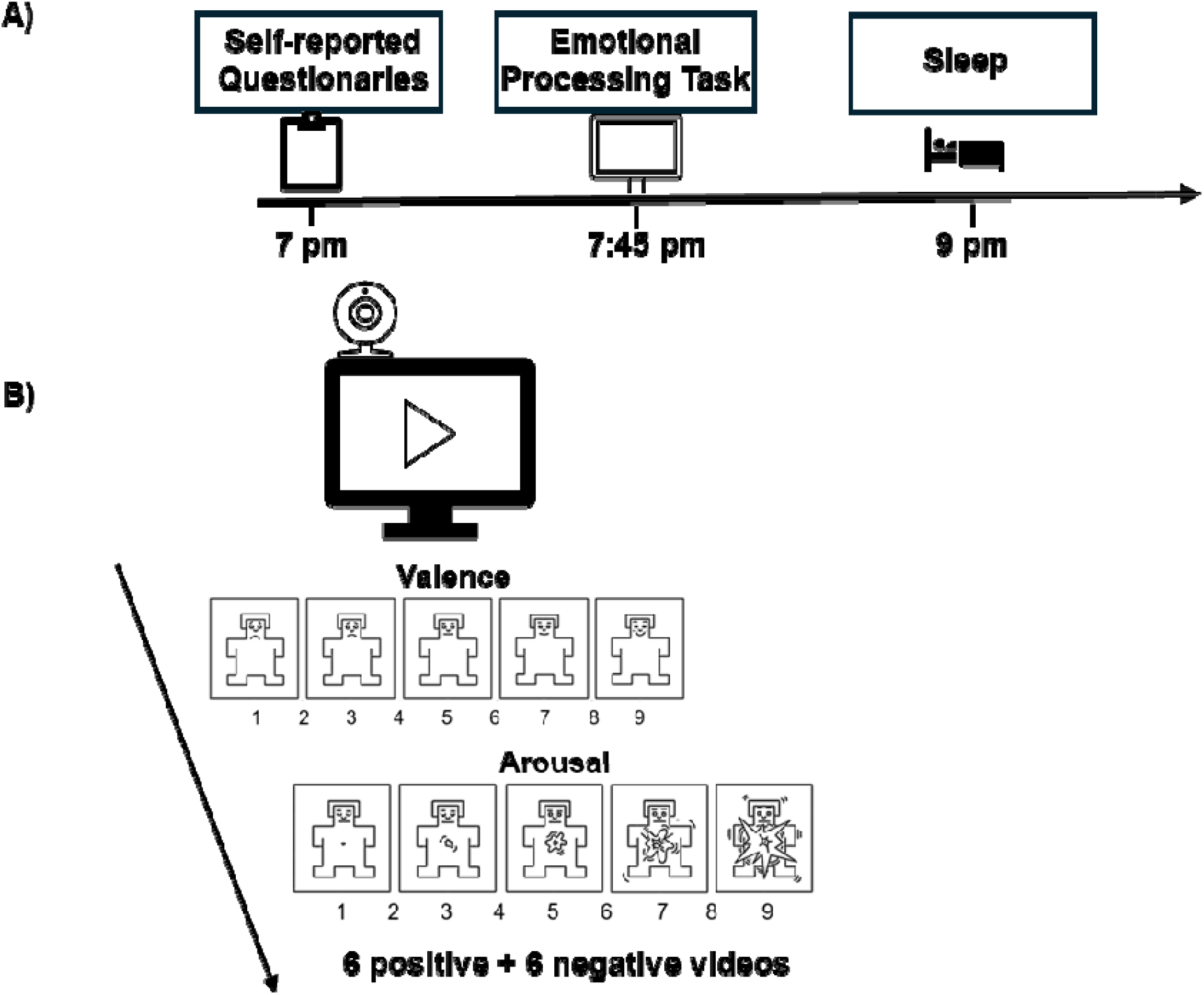
Study Design. **(A)** Participants arrived at the laboratory approximately 2 hours prior to their habitual bedtime and completed self-report questionnaires and an emotional regulation task before overnight polysomnography (PSG) recording. The figure depicts a hypothetical participant with a habitual bedtime of 9 pm. **(B)** During the emotional processing task, participants viewed 12 movie clips (6 positive, 6 negative), rated the valence and arousal of each clip using the Self-Assessment Manikin (SAM), and were given the option to look away from the screen if they felt uncomfortable. Facial expressions and look-away behavior were recorded via webcam.

#### Polysomnography (PSG)

Sleep was monitored with a 32 channel, Bluetooth-based EEG acquisition system with a sampling rate of 500 Hz (Live Amp 32, Brain Vision, NC) and EEG caps (Easy Cap GmbH, Herrsching, Germany) with 30 active electrodes for EEG, 2 electrodes for electrooculography (EOG), and 3 external electrodes for submental electromyography (EMG). Impedances were kept < 15 kOhm during set up.

The EEG data were bandpass filtered for 0.3-35 Hz and notch filtered at 60 Hz. An expert sleep scorer visually classified each 30-second epoch as N1, N2, N3, REM, or wake based on standard scoring criteria^55^. The measures of interest for sleep architecture were: total time in bed, total sleep time, wake after sleep onset (WASO, measured as the total minutes awake after sleep onset), sleep efficiency (calculated as the percentage of total sleep time relative to time in bed), sleep onset latency (the time from lights-off to the first occurrence of three consecutive sleep epochs), and the duration and percentage of each sleep stage.

Data for N2 and N3 sleep were preprocessed and analyzed with BrainVision Analyzer 2.0 (BrainProducts, Germany) and custom MATLAB (MathWorks, Natick MA) scripts. Data were visually examined by trained researchers blind to group assignment, and epochs containing artifacts were removed. Groups did not differ in the total duration of artifact-free N2 and N3 sleep (*F*(1,31), *p*=.21). Spherical spline interpolation was applied to EEG channels that displayed artifactual activity for more than 70% of the recording. Sleep spindles and SO were detected and characterized using previously validated automated algorithms^56^ and have been extensively described in earlier work^42,57–60^. A wavelet-based automated algorithm was used to detect slow (9.5-12.5 Hz) and fast (13.5-16.5 Hz) spindles in band-pass filtered data, using frequency ranges selected based on prior work in adolescent samples^61^. To enhance separation between spindle and non-spindle activity^62^, and consistent with earlier studies, slow and fast spindles were identified using an amplitude threshold set at nine times the median signal from artifact-free epochs and a minimum duration of 0.4 secs. We calculated the density (number of events per minute of artifact-free N2 and N3 sleep) of detected slow and fast spindles and characterized the morphological properties: slow/fast spindle duration (half-height width of the wavelet energy), slow/fast spindle frequency (spectral peak following Fast Fourier transform [FFT]), and slow/fast spindle amplitude (maximal voltage). For SO detection, the data were filtered between 0.5 and 4 Hz. Waveforms with amplitudes in the top 25% of each channel were used as the amplitude threshold, and the duration threshold was set at 0.8 secs. SO variables were as follows: SO density (number of detected SOs per minute of artifact-free N2 and N3 sleep), and SO peak-to-peak normalized amplitude.

#### Emotional Processing Task

The task, adapted from^63^ and programmed and presented using PsychoPy^64^, consisted of 12 age-appropriate movie clips depicting elation, joy, affection, fear, sadness, and disgust (2 videos per emotion) presented in a random order, ranging in duration from 82 to 186 sec. In each video, the positive or negative emotional content developed and intensified throughout. Participants were seated in front of a computer with a 13-inch monitor in a quiet testing room and were provided with noise-cancelling headphones to deliver the audio. Participants’ facial expressions were recorded throughout the task using a 720p/30fps webcam mounted on the computer screen. A poster showing a relaxing nature scene was placed on the wall to the participant’s right. Participants were informed that they would watch emotional movie clips and were instructed to look away from the screen toward the poster if they felt uncomfortable or upset continuing to watch the video. At the end of each trial participants rated the valence and arousal of the video clip using the Self-Assessment Manikin (SAM^65^) on a scale from 1 to 9. These subjective emotionality ratings were averaged separately for positive and negative movie clips.

Two trained raters, blind to group assignment and movie content, independently watched each webcam recording. Each video clip was divided into three equal segments. For each segment, raters coded the frequency and duration of look-away behavior, defined as any instance the participants turned away from the screen towards the poster. Look-away frequency and duration were then averaged across the three segments; thus, the resulting values represent the average frequency and duration per segment rather than totals across the entire video. Look-away behavior indexed behavioral emotion regulation. Raters also coded the valence and arousal intensity of the participants’ facial expressions using the SAM. Ratings were averaged across the three segments to obtain a score for each video clip. For facial expressions of valence and arousal, ratings were averaged when inter-rater differences were one or two points, for look-away frequency and duration, ratings were averaged when differences were ≤ 0.5. For discrepancies exceeding these thresholds, the raters jointly re-reviewed the relevant segment and reached a consensus rating. Inter-rater reliability, calculated from the independent ratings prior to consensus, was high (intraclass correlation coefficient [ICC]=.87, 95% CI [0.84, 0.90]). Facial expression ratings indexed participants’ expressive emotionality. All measures were averaged separately for positive and negative video clips.

#### Assessment of Youth Mental Health and Subjective Sleepiness

The revised Children’s Anxiety and Depression Scale-Parent Version (RCADS-P^66^) was used to assess parental reports of child depression and anxiety, which provides a depression subscore and two summary measures: total anxiety score (sum of separation anxiety, social phobia, generalized anxiety, panic disorder, obsessive-compulsive disorder subscales) and total score (sum of total anxiety score and the depression subscale). To complement the RCADS, parents also completed the Achenbach System of Empirically Based Assessments, Child Behavior Checklist (CBCL^67,68^), which assesses emotional and behavioral problems youth may experience. Age and sex normed internalizing and externalizing problem t-scores were utilized.

Additionally, participants completed the Positive and Negative Affect Schedule for Children (PANAS-C^69^), and scores were averaged for negative and positive affect items. The Confusion Hubbub and Order Scale (CHAOS^70^) was administered to evaluate the home environment structure. Participants also completed the self-reported Pubertal Development Scale^71^ and the Pediatric Daytime Sleepiness Scale (PDSS^72^).

### Statistical Analyses

#### Group comparisons

Parent and self-report scales, emotional processing task measures and sleep architecture variables were compared between groups using Welch’s independent samples t-tests, given unequal variances across most comparisons.

To examine group differences in NREM oscillations, independent samples t-tests were conducted at each electrode for each NREM oscillatory measure. Multiple comparisons across electrodes were addressed using a nonparametric cluster-based permutation method implemented in MATLAB^73^. Clusters consisted of neighboring electrodes that reached an uncorrected threshold of *p* < 0.05. Cluster statistics were defined as the sum of the test statistics across electrodes within each cluster. Permutation distributions were generated using 1000 iterations at each electrode, retaining the cluster with the maximum statistic from each permutation. Cluster-level corrected *p*-values reflected the proportion of permutation-derived maximum cluster statistics that exceeded the observed cluster statistic. To enhance replicability, and in line with EEG methodological recommendations^74^, we reported the highest effect size within each significant cluster (Cohen’s *d*).

#### NREM oscillation regression analyses

To examine associations of NREM oscillation measures with mood, affect and emotional processing task variables, separate linear regression analyses were performed at each electrode for each NREM oscillation measure and outcome. Each model included a single NREM oscillation predictor; spindle and SO measures were not entered simultaneously as predictors. The same cluster-based permutation method was used to correct for multiple comparisons. This procedure controlled for multiple comparisons across electrodes within each analysis; no additional correction was applied across the different clinical, affective, or behavioral outcomes examined. Effect sizes were calculated and reported as *f*^2^ at each electrode.

### Additional correlation analyses

Associations between emotional processing task measures and internalizing symptoms were examined using Pearson correlations. No covariates were included in these analyses.

#### Data exclusion and analysis-specific missingness

Four NC and three ANX participants were excluded from NREM oscillation analyses because the topographical distribution of oscillations could not be reliably detected due to excessive artifacts in several EEG sensors. Two NC participants were excluded from the sleep architecture analyses due to disruptions in their recordings caused by technical problems. Six NC (one only excluded for positive videos) and three ANX participants were excluded from the facial expression analyses, and three NC and two ANX participants were excluded from look-away analyses due to software errors or insufficient visibility of facial expressions in the recordings, which prevented reliable behavioral coding. Lastly, three NC and one ANX participants did not complete all CBCL items and were not assigned a total score. Distributions of sleep and behavioral variables were inspected using Q-Q plots and Shapiro-Wilk tests. Potentially extreme observations were identified using standardized scores and inspected for evidence of data or measurement error. No observations were excluded solely on the basis of these distributional checks. For variables showing substantial departures from normality, group differences were additionally examined using Wilcoxon rank-sum tests as sensitivity analyses. Consequently, the final sample size for the NREM oscillation group analyses was 36 (22 NC, 14 ANX). Analyses examining associations between NREM oscillations and internalizing symptoms included the same sample, except for CBCL analyses, which included 33 participants (20 NC, 13 ANX). Analyses examining associations between NREM oscillations and facial expressions included 32 participants (20 NC, 12 ANX) for negative videos and 31 participants (19 NC, 12 ANX) for positive videos. Analyses of look-away measures included 34 participants (21 NC, 13 ANX).

## Results

### Sample Characteristics

The ANX group exhibited higher parent-reported RCADS total, anxiety, and depression scores, as well as higher CBCL internalizing and total problems, compared to NC (Table 1). Groups did not differ in CBCL externalizing problems. Youth in the ANX group also reported higher negative affect on the PANAS-C. No group differences were observed in positive affect, pubertal status, daytime sleepiness, or home environment. Medications are listed in Supplemental Table 1. Briefly, 9 of participants in the ANX group were prescribed selective serotonin reuptake inhibitors (SSRIs), including 2 who were also prescribed the non-benzodiazapine anxiolytic buspirone. Based on sleep diaries completed for the night preceding the PSG recording, no group differences were observed in sleep duration (ANX=572.1±82.4 min, NC=573.9±61.9 min, *t*(38)=-0.1, *p*=.94) or number of awakenings (ANX=2.4±1.5, NC=2.1±1.3, *t*(41) =0.7, *p*=.52).

### Sleep EEG

Groups did not differ in sleep duration or architecture on the PSG night (Table 2). Average duration of slow spindles was significantly reduced in the ANX group (Figure 2, cluster size= 7 electrodes, *t_sum_*=-16.4, *p_corrected_*=.04; C3: *d*=0.9; effect sizes reflect the maximum value within each significant cluster). There were no other group differences in slow or fast spindle or SO measures (see Supplemental Figure 1 for topographical maps of NREM oscillations across groups). We also explored associations of age, sex, and pubertal status with sleep measures. Female participants had higher %N2 (*t(*38)=3.40, *p*=.002) and lower %N3 (*t*(38)=-4.40, *p*<.001), as well as lower spindle and SO amplitudes and marginally higher spindle density. Older age and more advanced pubertal status were associated with lower slow and fast spindle amplitudes. These exploratory analyses are detailed in Supplemental Figures 2 and 3.

**Figure 2.**
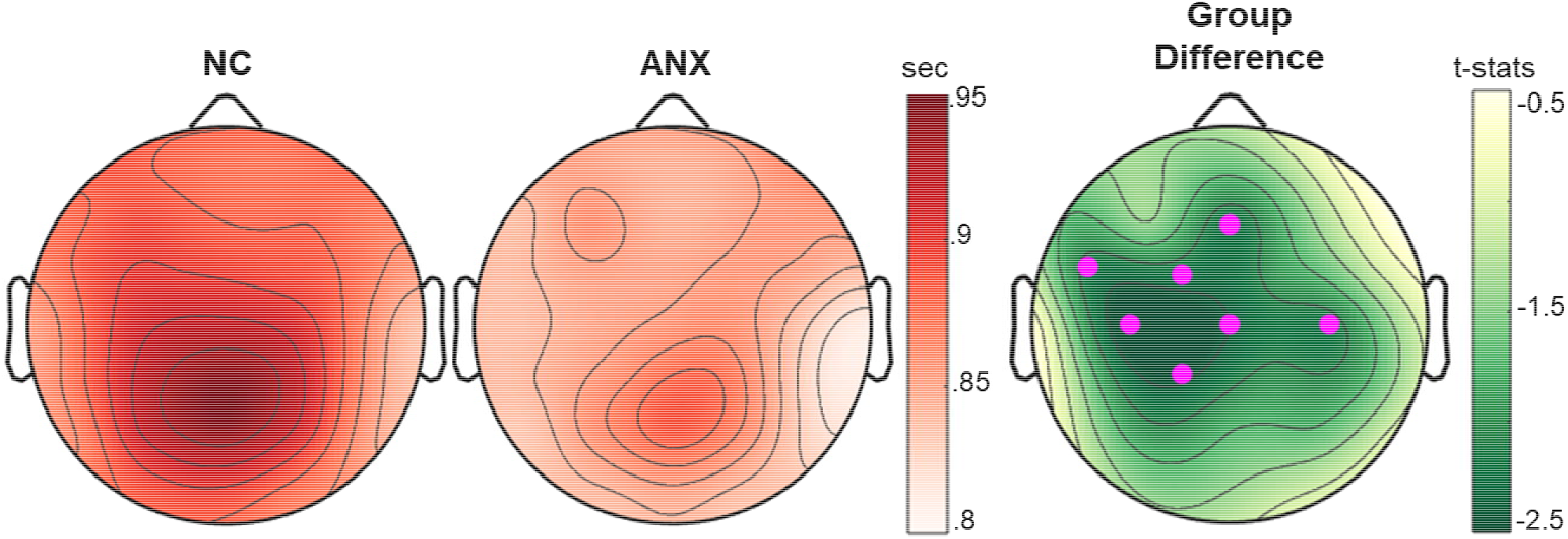
Group differences in slow spindle duration. Topographical maps of the average duration of detected slow sleep spindles with warmer colors representing longer duration in seconds. The topographical map on the right displays the t-statistics for the group comparison, with lighter colors indicating lower t-values. Electrodes marked in magenta surpass cluster-level correction for multiple comparisons (*p* _corrected_ < 0.05).

**Table 2.** Sleep quality and architecture.

|  | Anxiety<br>(n=16) |  | Comparison<br>(n=24) |  | *t* | *p* | Cohen's *d* | 95% CI |
| --- | --- | --- | --- | --- | --- | --- | --- | --- |
|  | *m* | *SD* | *m* | *SD* |  |  |  |  |
| TIB<sup>a</sup> (min) | 524.3 | 79.8 | 538.5 | 87.7 | -0.5 | .60 | 0.2 | -0.8;0.5 |
| TST<sup>b</sup> (min) | 484.6 | 66.3 | 494.6 | 92.6 | -0.4 | .69 | 0.1 | -0.8;0.5 |
| SOL<sup>c</sup> (min) | 19.6 | 20.5 | 16.3 | 12.5 | 0.6 | .58 | 0.2 | -0.5;0.8 |
| WASO<sup>d</sup> (min) | 8.7 | 17.7 | 5.2 | 6.3 | 0.8 | .46 | 0.3 | -0.4;0.9 |
| Sleep Efficiency (%) | 93.0 | 6.8 | 92.2 | 11.0 | 0.3 | .80 | 0.1 | -0.6;0.7 |
| N1(%) | 3.0 | 3.2 | 2.5 | 2.3 | 0.5 | .60 | 0.2 | -0.5;0.8 |
| N2(%) | 45.8 | 8.3 | 49.6 | 9.8 | -1.3 | .20 | 0.4 | -1.1;0.2 |
| N3(%) | 27.3 | 8.0 | 27.1 | 10.2 | 0.1 | .95 | 0.02 | -0.6;0.7 |
| REM(%) | 23.9 | 8.2 | 20.8 | 5.1 | 1.4 | .19 | 0.5 | -0.2;1.1 |
| N1(min) | 14.5 | 15.5 | 12.9 | 11.9 | 0.4 | .73 | 0.1 | -0.5;0.8 |
| N2(min) | 222.5 | 57.0 | 243.0 | 55.3 | -1.1 | .27 | 0.4 | -1.0;0.3 |
| N3(min) | 129.6 | 33.0 | 133.0 | 51.2 | -0.3 | .80 | 0.1 | -0.7;0.6 |
| REM(min) | 116.4 | 44.8 | 105.7 | 38.0 | 0.8 | .44 | 0.3 | -0.4;0.9 |

<sup>a</sup>Time in Bed; <sup>b</sup>Total Sleep Time; <sup>c</sup>Sleep Onset Latency; <sup>d</sup>Wake After Sleep Onset.

### Relations of NREM oscillations with mental health

We observed significant associations between spindle duration and internalizing symptoms across participants. Shorter slow spindle duration was associated with higher RCADS total scores (Figure 3A, 20 electrodes, *t_sum_*=-58.9, *p_corrected_*=.002; C3: *f*^2^=0.5). Exploratory analyses indicated this association was evident for both anxiety (Supplementary Figure 4A, 18 electrodes, *t_sum_*=-51.1, *p_corrected_*=.002; CP1: *f*^2^=0.4) and depression symptoms (Supplementary Figure 4B, 24 electrodes, *t_sum_*=-75.8, *p_corrected_*<.001; C3: *f*^2^=0.8). A similar pattern was observed for CBCL internalizing problems (Supplementary Figure 4C, 15 electrodes, *t_sum_*=-41.7, *p_corrected_*=.01; FC1: *f*^2^=0.4), whereas there was no association between spindle duration and CBCL externalizing problems. Fast spindle duration was also negatively associated with internalizing symptoms, although the effect was weaker (RCADS total score, Supplementary Figure 4D, 6 electrodes, *t_sum_*=-13.5, *p_corrected_*=.04; Fp2: *f*^2^=0.2).

**Figure 3.**
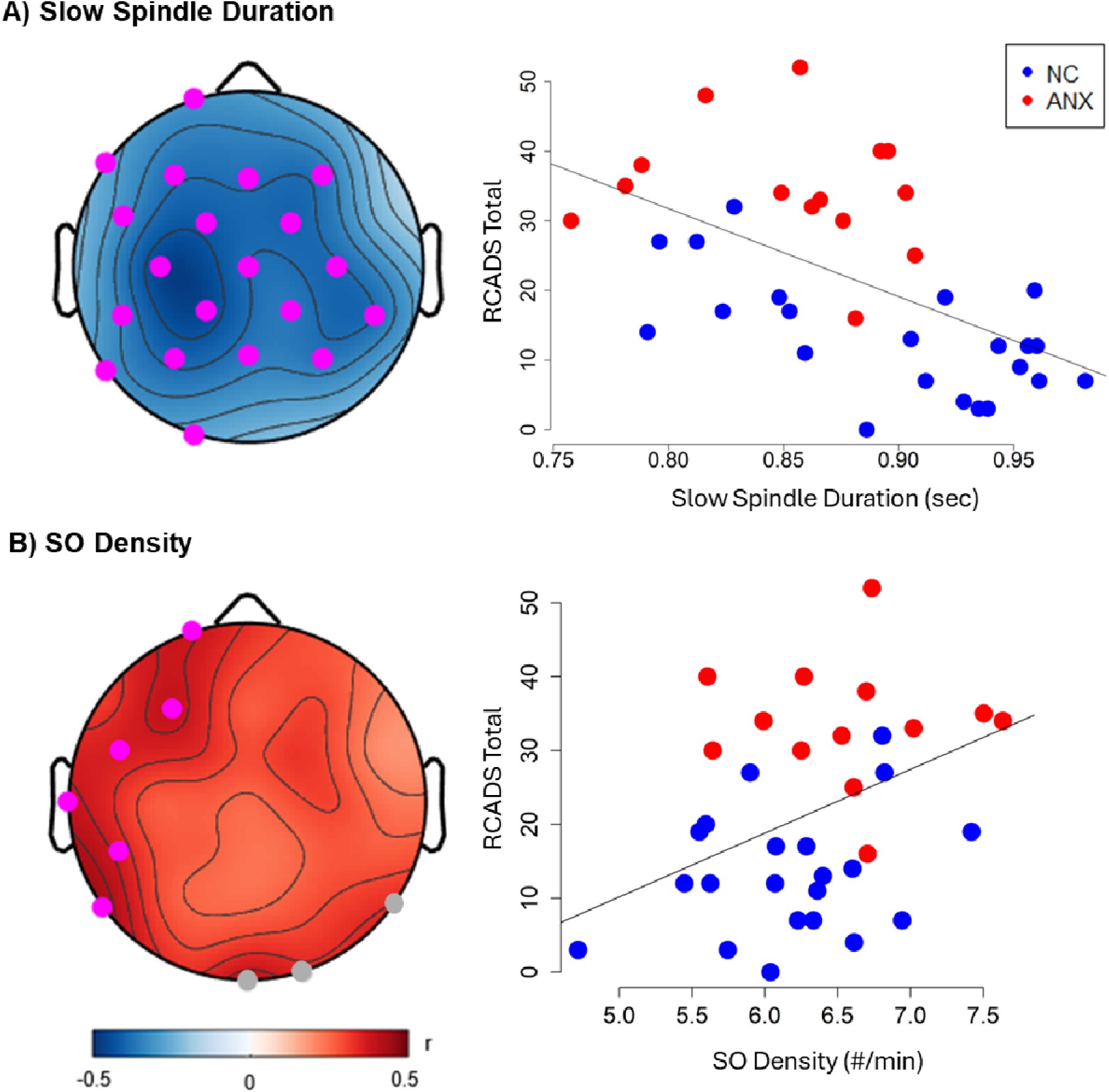
Associations of NREM Oscillations with Internalizing Symptoms. Parents completed the Revised Child Anxiety and Depression Scale (RCADS). The Total Score was used as a measure of internalizing symptom severity. **(A)** Slow spindle duration was negatively associated with RCADS total score, whereas **(B)** SO density was positively associated with RCADS total score. Left panels show topographical maps of correlation strength (Pearson’s r). NREM oscillation measures shown in the figures represent averages across significant channels. Warm colors represent positive correlations and blue represents negative correlations. Electrodes marked in gray correspond to p_uncorrected_<.05 and electrodes in magenta surpass cluster-level correction for multiple comparisons (p_corrected_ < .05). The right panels show scatterplots of these relationships in the Comparison (blue) and Anxiety (red) groups. The regression line is shown for the entire sample.

Extending the spindle findings, SO density was positively associated with internalizing symptoms. Across the entire sample, higher SO density was associated with higher RCADS total score (Figure 3B, 6 electrodes; *t_sum_* = 14.6, *p_corrected_*= .05; P7: *f²* = 0.2). This association was stronger for depressive symptoms (Supplementary Figure 5A; 25 electrodes; *t_sum_* = 61.7, *p_corrected_*= .01; P7: *f²* = 0.3) than for anxiety symptoms, which did not reach significance (Supplementary Figure 5B; 6 electrodes; *t_sum_* = 13.6, *p_corrected_*= .06; P7: *f²* = 0.2). A similar pattern was observed for CBCL internalizing problems (Supplementary Figure 5C; 22 electrodes; *t_sum_* = 52.3, *p_corrected_*= .01; P7: *f²* = 0.3). Because slow oscillatory activity is sensitive to homeostatic sleep pressure, we also examined whether self-reported sleep duration on the night prior to PSG was associated with SO density. Regression coefficients and standard errors for significant associations are provided in Supplementary Table 2. No significant association was observed (*p* = .49).

To investigate whether SO density and slow spindle duration captured overlapping aspects of NREM physiology, we first examined the relationship between the two significant clusters predicting RCADS Total score. SO density and slow spindle duration were significantly and negatively correlated (*r*=-.55, *p*<.001), indicating that shorter slow spindle duration was associated with higher SO density. We then entered both cluster measures into the same regression model predicting RCADS total score and assessed multicollinearity using variance inflation factors (VIF). Both predictors had VIF values of 1.4, indicating minimal multicollinearity. In summary, both shorter spindle duration and higher SO density were associated with higher internalizing symptoms across the entire sample of young adolescents. We did not observe associations between other spindle or SO measures and mental health variables.

### Emotional Processing Task

Task measures are summarized in Table 3. The groups did not differ in self-ratings of valence or arousal for positive or negative movie clips. Youth in the ANX group looked away from the screen more frequently and for longer durations than NC during positive, but not negative videos. The arousal intensity of facial expressions during negative movie clips was higher in the ANX group, with no other group differences observed (all *p*’s > .07).

**Table 3.** Emotional Processing Task.

|  | Anxiety<br>(n=14) |  | Comparison<br>(n=20) |  | *t* | *p* | Cohen's *d* | 95% CI |
| --- | --- | --- | --- | --- | --- | --- | --- | --- |
|  | *m* | *SD* | *m* | *SD* |  |  |  |  |
| *Self-reported emotionality ratings for movie clips<sup>a</sup>* |  |  |  |  |  |  |  |  |
| Valence (positive) | 7.3 | 0.5 | 7.2 | 0.6 | 0.9 | .39 | 0.3 | -0.4;0.9 |
| Valence (negative) | 2.8 | 1.2 | 3.1 | 0.9 | -1.0 | .33 | 0.3 | -1.0;0.3 |
| Arousal (positive) | 5.5 | 1.3 | 5.6 | 1.1 | -0.3 | .80 | 0.1 | -0.7;0.6 |
| Arousal (negative) | 6.1 | 1.5 | 5.8 | 1.3 | 0.7 | .46 | 0.2 | -0.4;0.9 |
| *Facial expressions of emotions<sup>b</sup>* |  |  |  |  |  |  |  |  |
| Valence (positive) | 5.3 | 1 | 5.0 | 0.7 | 0.7 | .47 | 0.3 | -0.4;1.0 |
| Valence (negative) | 4.3 | 0.7 | 4.6 | 0.6 | -1.4 | .16 | 0.5 | -1.2;0.2 |
| Arousal (positive) | 3.4 | 1.6 | 2.6 | 1.3 | 1.5 | .14 | 0.5 | -0.2;1.2 |
| Arousal (negative) | 4.2 | 1.3 | 2.7 | 1.1 | 3.5 | .002* | 1.2 | 0.5;2.0 |
| *Look-away events<sup>c</sup>* |  |  |  |  |  |  |  |  |
| mean # (positive) | 1.9 | 1.6 | 0.8 | 0.6 | 2.4 | .03* | 0.9 | 0.1;1.6 |
| mean # (negative) | 1.2 | 0.9 | 0.8 | 0.5 | 1.9 | .07 | 0.7 | -0.03;1.3 |
| duration (sec; positive) | 2.4 | 2.3 | 0.9 | 1.0 | 2.3 | .04* | 0.8 | 0.1;1.5 |
| duration (sec; negative) | 2.3 | 2.9 | 1.9 | 2.3 | 0.5 | .62 | 0.2 | -0.5;0.8 |

<sup>a</sup>Participants rated the valence and arousal of each movie clip using the Self-Assessment Manikin (SAM<sup>65</sup>) on a scale from 1 to 9
<sup>b</sup>Facial expressions of the participants while watching the movie clips were rated on SAM scales by two researchers independently;
<sup>c</sup>Participants were instructed to look away from the screen if they felt uncomfortable while watching the videos. Values represent average frequency and duration per segment across the three equally divided video segments.
\**p*<.05

The intensity of facial expression arousal during negative videos was associated with higher internalizing symptoms. Correlation analyses showed that higher internalizing symptoms were significantly associated with greater facial expression of arousal across measures (e.g. CBCL internalizing t-scores: *r*=.5, *p*=.003; RCADS total:*r*=.5, *p*=.01). Look-away behavior was positively associated with internalizing behaviors such that youth with higher depression and anxiety symptoms looked away from both the positive and negative movie clips more frequently and for longer durations (e.g. RCADS total, frequency of look away events for negative videos: *r*=.5, *p*=.004; positive videos: *r*=.4, *p*=.01). Children’s own ratings of arousal for the negative videos were marginally related to RCADS depression subscores (*r*=.3, *p*=.05); indicating that children who rate negative videos as more arousing had higher depression symptoms. Other self-ratings of valence or arousal of these movie clips were not associated with mental health variables (all *p*’s >.08).

### Relations of NREM Oscillations with Emotional Processing Task

To examine whether behavioral and expressive indices of emotion while viewing videos were associated with NREM oscillations during subsequent sleep, we tested associations of facial arousal and look-away behaviors, separately for positive and negative videos, with spindle and SO measures. For the negative videos, both the intensity of facial arousal (Figure 4A, 7 electrodes, *t_sum_*=-17.4, *p_corrected_*=.04; FC1: *f* ^2^=0.3), and frequency of looking away from the screen (Figure 4B, 9 electrodes, *t_sum_*=-20.1, *p_corrected_*=.03; F3: *f* ^2^=0.2) were negatively associated with slow spindle duration, such that youth who expressed greater facial arousal and looked away from the screen more frequently during negative movie clips had shorter spindle duration during subsequent sleep. No other associations were significant.

**Figure 4.**
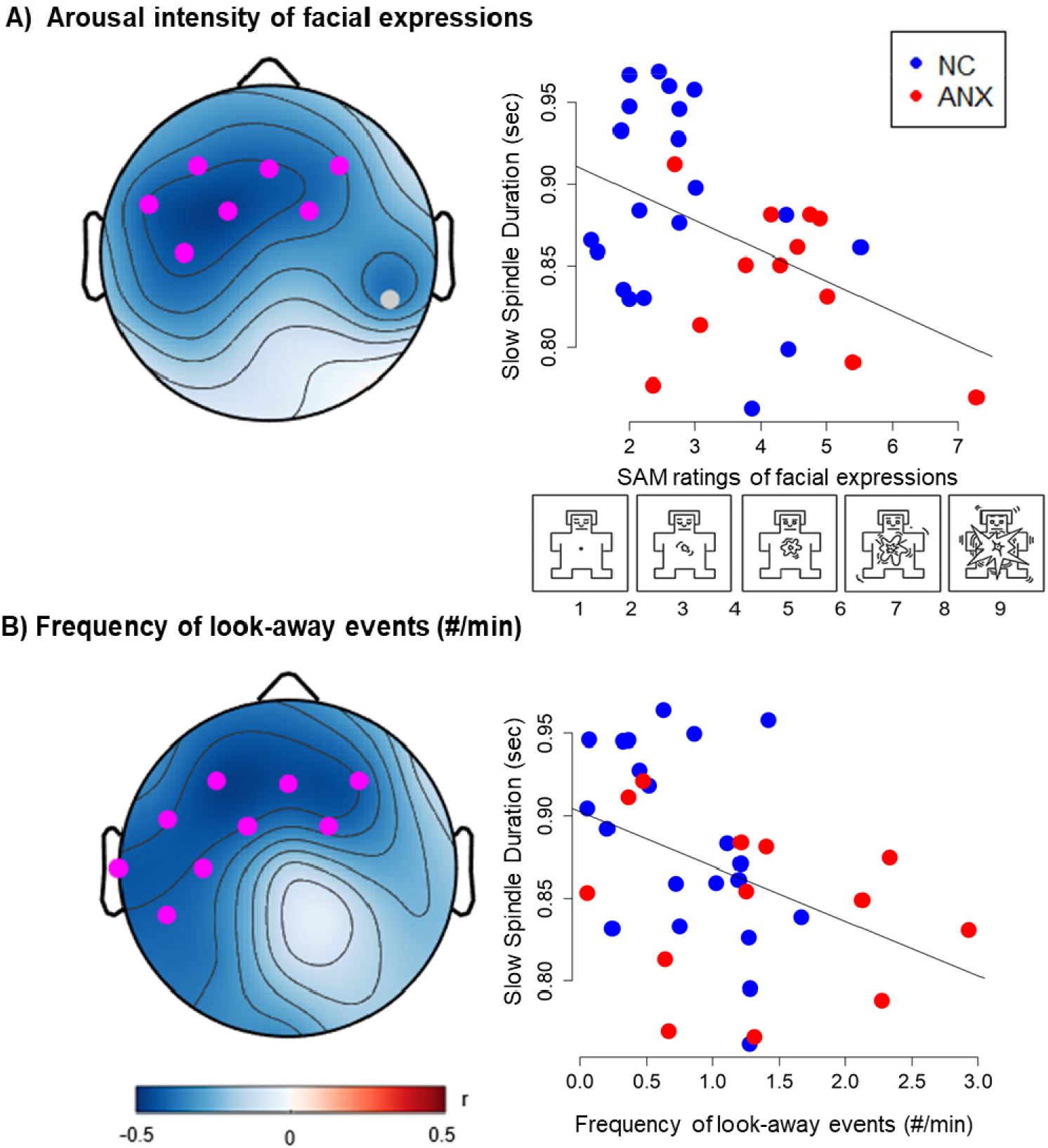
Associations of Slow Spindle Duration with Emotional Processing Task: **(A)** Arousal intensity of facial expressions during negative movie clips (higher ratings= more arousing facial expressions; ratings averaged across all negative movie clips) was associated negatively with slow spindle duration during subsequent sleep**, (B)** Participants were instructed to look away from the screen to a relaxing nature themed poster on the wall if they felt uncomfortable while watching the movie clips. The average frequency of look-away events during negative movie clips was negatively associated with slow spindle duration during subsequent sleep. Left panels show topographical maps of correlation strength (Pearson’s r). Warm colors represent positive correlations and blue represents negative correlations. Electrodes marked in gray correspond to p_uncorrected_<.05 and electrodes in magenta surpass cluster-level correction for multiple comparisons (p_corrected_ < .05). The right panels show scatterplots of these relationships in the Comparison (blue) and Anxiety (red) groups. The regression line is shown for the entire sample.

## Discussion

In this study, we characterized NREM sleep physiology and emotional functioning in children with anxiety disorders and a demographically matched comparison group during late childhood and early adolescence, a developmental period marked by substantial changes in both sleep neurophysiology and vulnerability to internalizing psychopathology. While sleep duration and architecture did not differ between groups, we observed evidence for alterations in NREM sleep microarchitecture associated with internalizing symptoms and emotional behavior. Specifically, slow spindle duration was shorter in the ANX group and associated with higher anxiety and depression symptoms across participants, whereas greater SO density was associated primarily with depressive symptoms. In addition, greater facial emotional arousal and more frequent look-away behavior (i.e. avoidance) during negative movie clips were associated both with higher internalizing symptoms and with shorter slow spindle duration during subsequent sleep. Together, these findings suggest that specific features of NREM sleep microarchitecture, particularly spindle duration, may be relevant to emotional functioning and internalizing symptoms during early adolescence.

The observed alterations in NREM oscillatory activity may reflect broader disruptions in thalamocortical processes implicated in the development of psychopathology during adolescence. Specifically, we found reduced slow spindle duration in youth with anxiety disorders in the absence of differences in overall sleep quality or architecture. These findings highlight that alterations in sleep microarchitecture may be detectable even in the absence of differences in conventional measures of sleep macrostructure. The group difference in spindle duration partially aligns with prior work in older adolescents with social anxiety disorder by Wilhelm and colleagues^53^, who reported widespread reductions in sigma-band activity, particularly in the 13-16 Hz range, using spectral power analyses rather than spindle detection. Importantly, they also reported links between spindle activity and severity of anxiety symptoms and self-reported arousal ratings for negative picture stimuli viewed prior to sleep, supporting the broader idea that spindle abnormalities may relate to affective dysregulation during adolescence. In contrast, a high-density EEG study of adolescents and young adults at clinical high risk for psychosis reported reduced spindle duration and interpreted these alterations as a potential early biomarker of psychosis risk and associated cognitive dysfunction^75^. Together, these findings suggest that alterations in spindle activity may emerge across adolescent psychopathology, although the specific spindle features affected and their functional implications may differ across diagnostic groups and developmental stages.

Beyond categorical group differences, shorter spindle duration was associated dimensionally with greater internalizing symptoms and heightened emotional responding across participants in our sample. Specifically, youth with shorter spindle duration exhibited higher anxiety and depressive symptom severity, greater facial emotional arousal, and more frequent look-away behavior while viewing negative movie clips. These findings extend prior work linking sleep spindles to affective and cognitive functioning^33,76–78^ and suggest that spindle duration may index individual differences in affective processes and regulatory strategies during adolescence. Although relatively few studies have specifically examined spindle duration, prior work suggests this parameter may relate to broader cognitive and affective functioning. For example, a meta-analysis reported modest positive associations between slow spindle duration and general cognitive ability, although effects were weaker and less consistent than those observed for spindle amplitude^79^. Within this context, the present findings raise the possibility that shorter spindle duration may reflect altered sleep-dependent processes involved in emotional processing during a sensitive developmental period. Notably, these associations were evident across both anxiety and depressive symptoms, consistent with the findings of substantial overlap between these forms of internalizing problems during adolescent development^80^.

Findings relating sleep spindles to anxiety and internalizing symptoms have varied across studies, although several methodological and conceptual differences may help contextualize these discrepancies. For example, Meers and colleagues^52^ did not observe group differences in spindle activity between prepubertal children with GAD and comparison subjects but reported positive associations between spindle density and worry severity across the entire sample. In contrast, in a non-psychiatric sample of prepubertal children, the same group reported that greater spindle density was associated with lower arousal to negative affective stimuli and greater use of adaptive positive emotion regulation strategies^76^, supporting a broader relationship between spindle activity and emotional functioning rather than categorical psychopathology. Similarly, reduced spindle density has also been reported in youth with major depressive disorders and those at familial high risk for depression^49^. Together with the present findings, this literature suggests that distinct spindle features may differentially relate to dimensions of emotional processing and internalizing symptoms. Developmental stage may also be particularly important as spindle characteristics undergo substantial maturational change during adolescence^81^, potentially contributing to variability across studies in both the direction and magnitude of observed associations.

Notably, most associations in the present study were observed for slow rather than fast spindles. Slow and fast spindles differ in their topographic distributions, with slow spindles showing a more frontal predominance and fast spindles a more centroparietal distribution^82^. They also appear to serve partially distinct functions^83^, with fast spindles being more consistently linked to memory consolidation^59,84–86^. However, relatively little is known about potential differences between spindle subtypes in emotional functioning. The predominance of slow spindle findings in the present study may suggest that slow spindles are particularly relevant to affective processes and internalizing symptoms, although future studies are needed to directly test this possibility.

In contrast to the spindle findings, greater SO density was positively associated with internalizing symptoms, particularly with depression. Notably, SO density was negatively correlated with slow spindle duration, indicating that youth with shorter spindle duration tended to exhibit higher SO density. Slow oscillatory activity has been less frequently examined in relation to internalizing psychopathology. Prior studies in non-psychiatric samples of young adults have linked SO and slow wave activity (SWA; delta power) with lower anxiety and negative affect upon awakening^38,39^. These findings appear to contrast with the present results. One possibility is that the functional significance of SO activity may differ across development. During puberty, SO activity undergoes substantial maturational change, which has been proposed to reflect ongoing cortical development, changes in synaptic strength and plasticity-related processes^87,88^. Consequently, greater SO density in youth with elevated internalizing symptoms may not necessarily reflect more efficient sleep-dependent emotional regulation but instead may index individual differences in neurodevelopmental processes. Although speculative, this interpretation is consistent with models linking SO to cortical maturation and the emergence of psychopathology during adolescence^89^. Notably, SO density was negatively correlated with slow spindle duration, indicating that youth with shorter spindle duration tended to exhibit higher SO density. The moderate correlation between SO density and spindle duration suggests that these measures may reflect partially overlapping aspects of NREM sleep physiology.

Youth with anxiety disorders exhibited greater facial emotional arousal during negative movie clips and more frequent look-away behavior during the task, suggesting heightened emotional responding to affective stimuli. Importantly, both facial emotional arousal and look-away behavior were associated with internalizing symptoms across the entire sample, indicating that these behavioral indices tracked broader variation in affective functioning. The emotional processing task occurred immediately before sleep, whereas much of the prior literature has examined NREM oscillations in relation to emotional memory consolidation or emotional processing assessed after sleep. This temporal proximity may have contributed to the observed associations with subsequent spindle characteristics; however, because the task was not experimentally manipulated, we cannot determine whether pre-sleep emotional responding influenced subsequent NREM physiology or whether these associations reflect more stable individual differences. Although look-away behavior was introduced to participants as an optional strategy to avoid discomfort while viewing emotional stimuli, youth with anxiety disorders demonstrated more look-away behavior during positive movie clips, and greater look-away behavior was correlated with higher internalizing symptoms, suggesting that this behavior may reflect a broader tendency to disengage from emotionally salient information, perhaps in service of reducing levels of arousal. However, because participants were not asked to report the reason for each look away instance, some may have reflected factors unrelated to emotional processing, such as distraction or attention lapses Further, no group differences or associations with internalizing symptoms or NREM oscillations were observed for self-reported SAM ratings. One possibility is that these measures capture subjective emotional experience, whereas facial expressions reflect directly observable and more automatic emotional expression, and therefor may capture individual differences not reflected in self-report in children. Lastly, look-away behavior was relatively infrequent, resulting in a restricted range that may have limited our ability to detect individual differences and associations with other measures.

This study should be considered in the context of several limitations. First, the sample size was modest, and data loss across some measures further reduced the available sample for specific analyses, limiting statistical power and the precision of effect size estimates. Replication in larger and more diverse samples will be important for establishing the robustness and generalizability of these findings. Although electrode-wise analyses were corrected for multiple comparisons using cluster-based permutation testing, no additional correction was applied across the multiple clinical and behavioral outcomes examined, increasing the possibility of false-positive findings. In addition, a substantial proportion of youth with anxiety disorders were receiving SSRIs, which may have influenced symptom expression and sleep physiology. Although prior studies have reported decreased REM sleep and increased NREM sleep with SSRI use^90–92^, no group differences in sleep architecture were observed in the present study. Moreover, there are no well-established effects of SSRIs on sleep spindle activity. The cross-sectional design and correlational analyses also prevent conclusions about developmental trajectories or causal relationships between NREM oscillations, emotional processing, and internalizing symptoms. Another limitation of the present study is that sleep was assessed during a single night of polysomnography without an adaptation night. First-night effects can influence sleep duration, continuity, and architecture^93^. Although efforts were made to minimize these effects by familiarizing participants with the laboratory beforehand, allowing a parent to stay in an adjacent room, encouraging familiar items from home, and conducting recordings in a bedroom-like environment, some first-night effects may have remained. Notably, groups did not differ in PSG-measured sleep duration, sleep efficiency, sleep onset latency, WASO, or sleep architecture, providing some reassurance that overall adaptation to the laboratory environment was comparable. Nevertheless, differential effects on NREM microarchitecture cannot be ruled out. Future studies incorporating an adaptation night will be important for establishing the robustness of the present findings. A further limitation is that youth with diagnosed sleep disorders were excluded; accordingly, these findings may not generalize to youth with anxiety and comorbid diagnosed sleep disorders. Finally, given the substantial maturational changes in sleep oscillations occurring during late childhood and adolescence, longitudinal studies spanning this developmental period will be important for clarifying whether alterations in spindle and SO activity reflect stable markers of internalizing psychopathology, developmental variation in thalamocortical maturation, or state-related changes associated with emotional functioning.

## Conclusion

This study provides evidence that specific features of NREM sleep microarchitecture are associated with emotional functioning and internalizing symptoms during late childhood and early adolescence, a developmental period marked by substantial changes in both sleep neurophysiology and vulnerability to psychopathology. Shorter spindle duration was observed in youth with anxiety disorders and was associated dimensionally with greater anxiety, depression, and internalizing symptoms, as well as heightened emotional responding to negative stimuli. In contrast, greater SO density was positively associated with internalizing symptoms, suggesting that distinct features of NREM oscillations may relate differently to affective processes during development. Importantly, these associations were observed in the absence of group differences in sleep duration or architecture, highlighting the value of examining sleep microarchitecture when investigating links between sleep and youth psychopathology. Together, these findings support the potential relevance of NREM sleep oscillations to emotional functioning during adolescence and underscore the importance of developmental approaches for understanding sleep-related mechanisms associated with internalizing psychopathology.

## Acknowledgments

This work was supported by an Iowa Neuroscience Institute Ideas Lab award to BB. Hazal Arpaci is the recipient of fellowship support through the National Institute of General Medical Sciences predoctoral training grants T32GM108540 and T32GM149386 (MPIs: Lutgendorf/Voss/Tranel). The authors would like to acknowledge the contributions of Josie Norris, Chani Parrott, Abby McLeod, Alexia Lynn, Avery Maeder, Ellie Ronnebaum, Fay Lin, and Meredith Sloat to data collection and curation. The authors thank the children and families who participated in the study.

## Author Contributions

Hazal Arpaci (Methodology, Software, Formal Analysis, Investigation, Data Curation, Visualization, Writing-original draft), Morgan Lott (Investigation, Data Curation), Anna Holowicki (Investigation), Sarah Dickens (Investigation, Data Curation), Aditi Tripathy(Investigation, Data Curation), Nandita Banik (Methodology, Investigation),Intisar Becic( Investigation), Narges Firouzshahi (Methodology), Candice Alfano (Conceptualization, Methodology, Writing-review & editing), Gerta Bardoshi (Conceptualization, Funding acquisition, Writing-review & editing), Bengi Baran (Conceptualization, Funding acquisition, Investigation, Methodology, Project administration, Formal Analysis, Writing-review & editing)

## Data Availability Statement

De-identified data are available from the authors upon reasonable request

