## Supplementary Information for "NREM sleep microarchitecture is associated with internalizing symptoms and emotional functioning in early adolescence"

**Supplemental Figure 1.** Topographical maps of group averages of the features of **A)** Slow Spindles; **B)** Fast Spindles; and **C)** Slow Oscillations. Warm colours represent higher values. Electrodes marked in grey correspond to *p*_uncorrected_<.05 and electrodes in magenta surpass cluster-level correction for multiple comparisons (*p*_corrected_ < .05). Only slow spindle duration was significantly different between groups after cluster corrections**.**

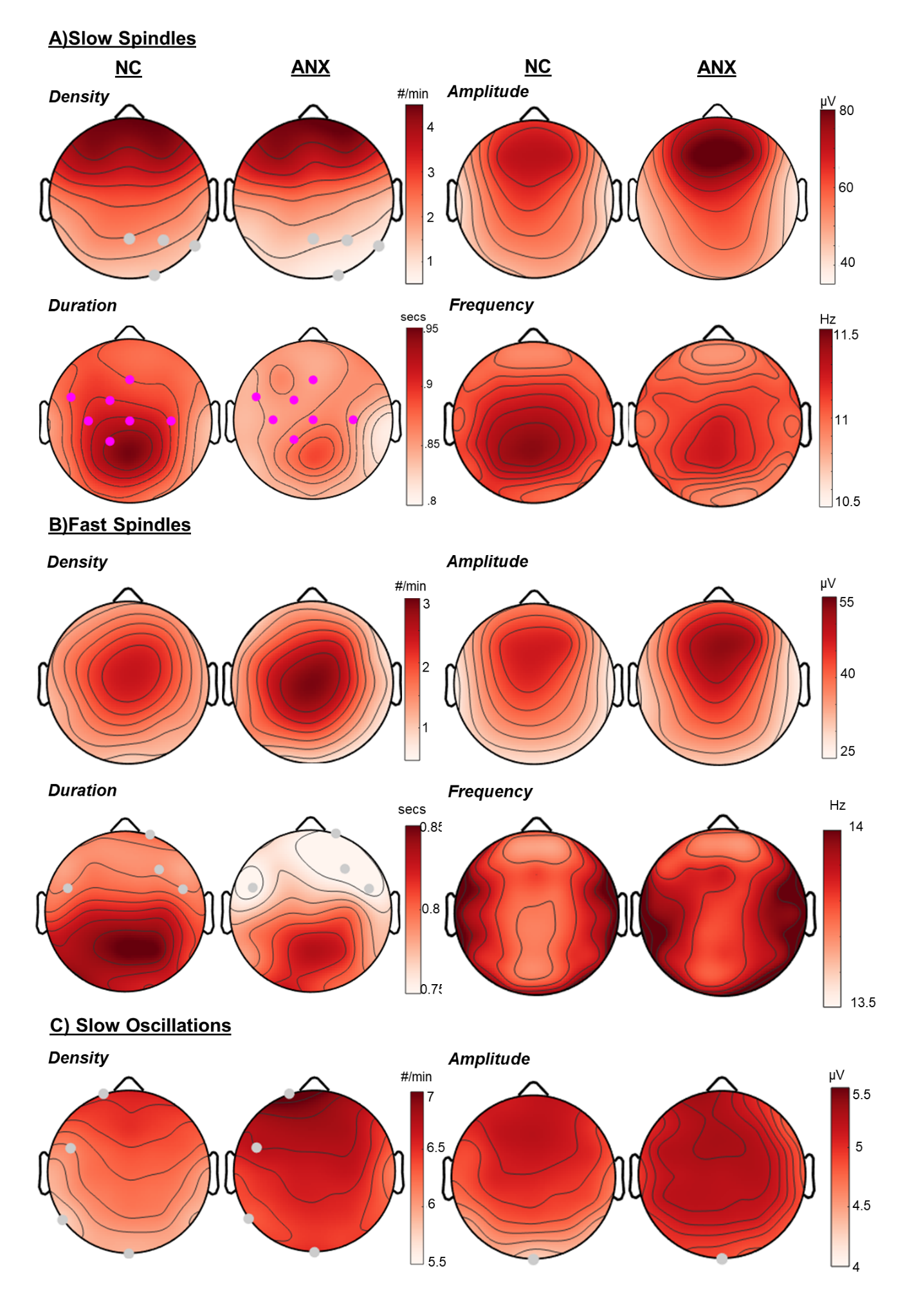

**Supplemental Figure 2.** Topographical maps of exploratory associations between NREM oscillatory features and age and pubertal status. **A)** Slow spindle amplitude was negatively associated with age (25 electrodes, *t_sum_*=-67.65, *p_corrected_* =.002); **B)** Fast spindle amplitude was negatively associated with age (7 electrodes, *t_sum_*=-15.02, *p_corrected_* = .05). **C)** Pubertal status (i.e. total score on Pubertal Development Scale) was negatively associated with slow spindle amplitude (16 electrodes, *t_sum_*=-38.69, *p_corrected_* = .03). Warm colours represent positive values and blue shading represents negative values. Electrodes marked in grey correspond to *p*_uncorrected_<.05 and electrodes in magenta surpass cluster-level correction for multiple comparisons (*p*_corrected_ < .05). The right panels show scatterplots of these relationships in the NC (blue) and ANX (red) groups. NREM oscillation measures shown in the scatterplots represent averages across significant electrodes.The regression line is shown for the entire sample.

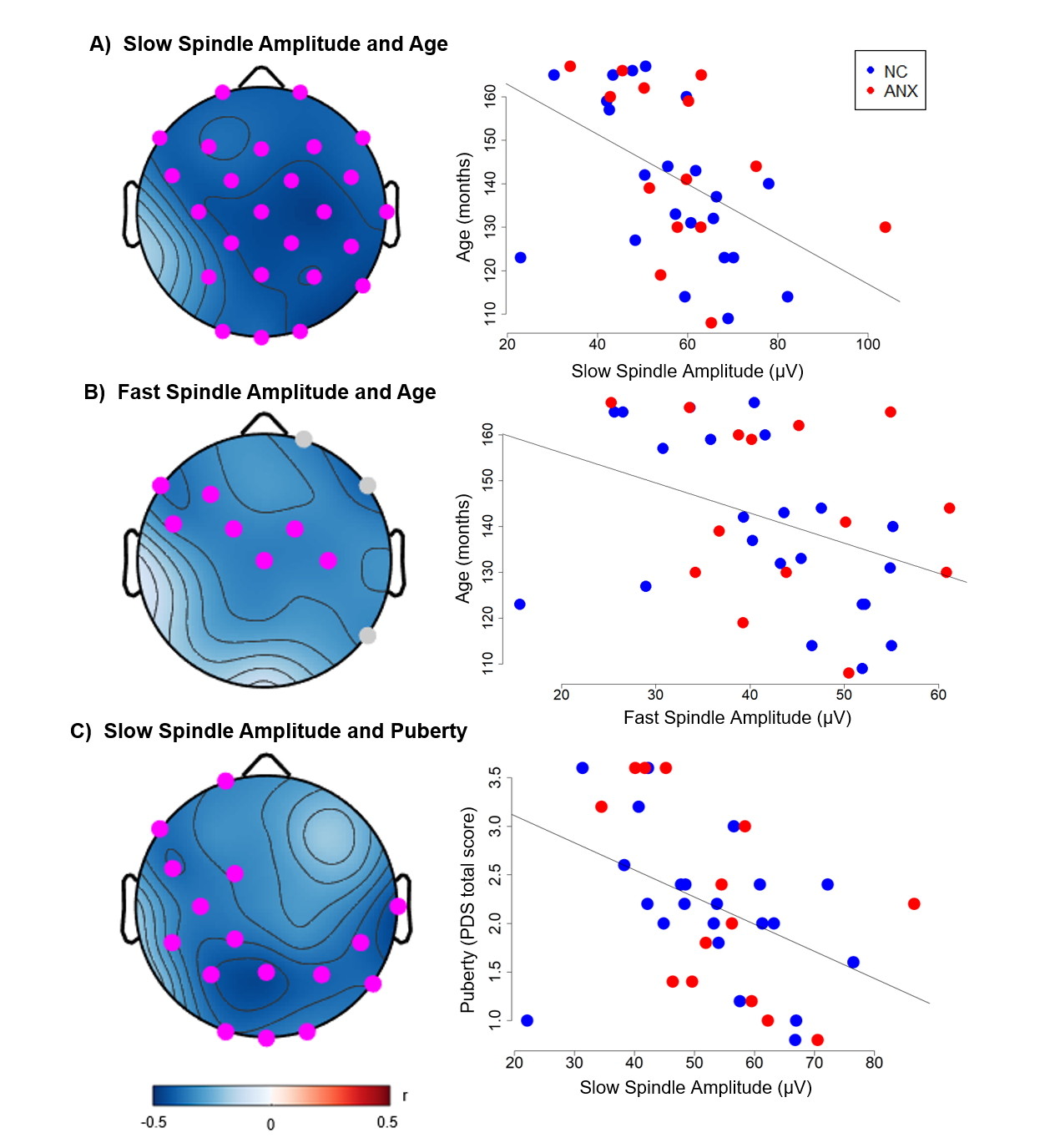

**Supplemental Figure 3.** Topographical maps of exploratory analyses examining associations between NREM oscillatory features and sex. Males had higher **A)** Slow spindle amplitude (22 electrodes, *t_sum_*=61.99, *p_corrected_* = .01); **B)** fast spindle amplitude (15 electrodes, *t_sum_*=38.35, *p_corrected_* = .02), and **C)** SO amplitude (12 electrodes, *t_sum_*=26.54, *p_corrected_* =. 03). On the other hand, females had higher **D)** fast spindle density (6 electrodes, *t_sum_*=-15.90, *p_corrected_* = .04), and **E)** marginally higher slow spindle density (7 electrodes, *t_sum_*= -16.60, *p_corrected_* = .06). Warm colours represent positive values, and blue shading represents negative values. Electrodes marked in grey correspond to *p*_uncorrected_<.05 and electrodes in magenta surpass cluster-level correction for multiple comparisons (*p*_corrected_ < .05).

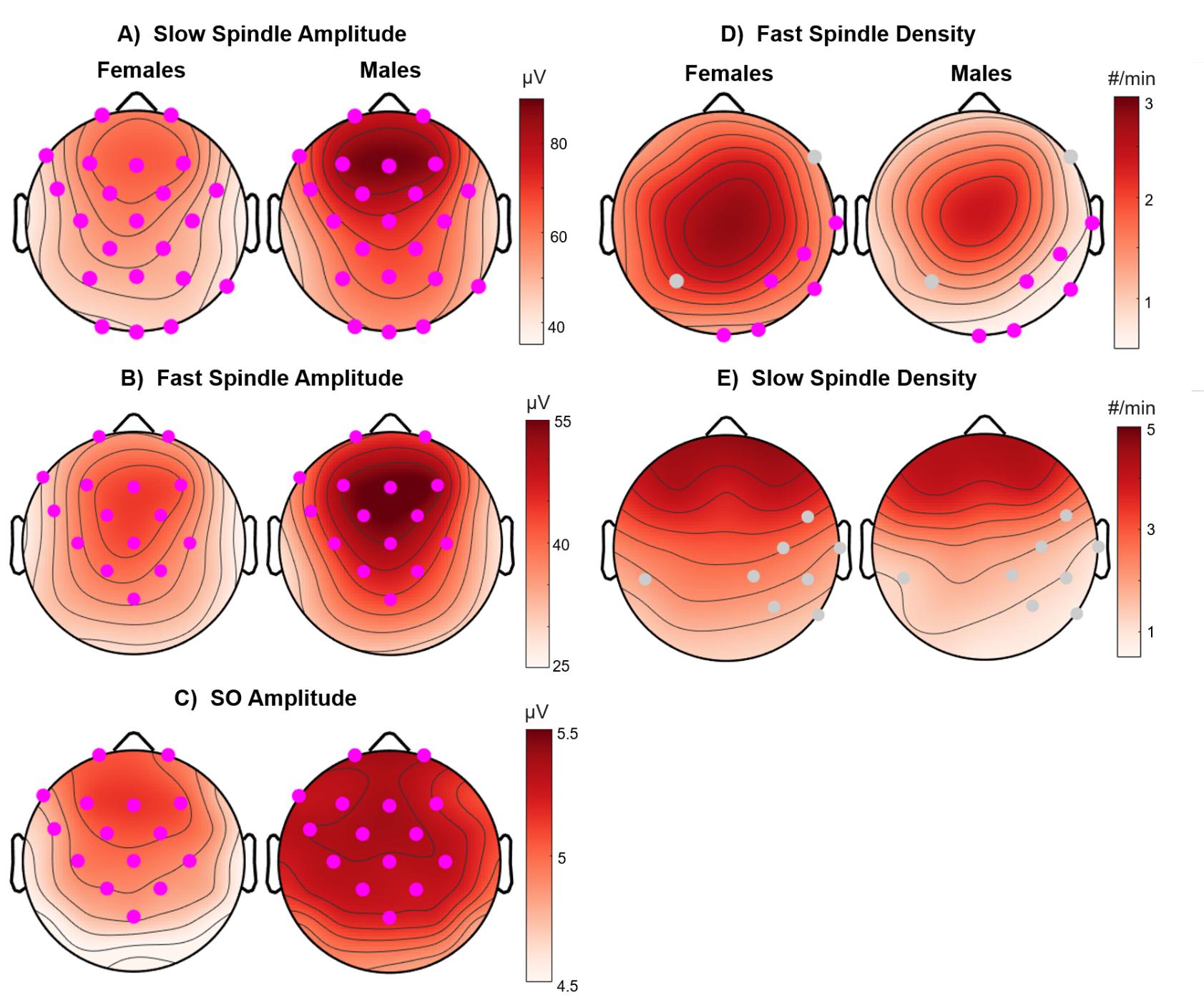

**Supplemental Figure 4. Association of NREM Spindle Duration with Internalizing Symptoms.** Parents completed the Revised Child Anxiety and Depression Scale (RCADS) and Child Behavior Checklist (CBCL). Slow spindle duration was negatively associated with (A) RCADS Anxiety Symptoms, (B) RCADS Depression Symptoms, and (C) CBCL Internalizing Problems. Similarly, (D) fast spindle duration was negatively associated with RCADS Total Score. Left panels show topographical maps of the correlation strength (Pearson's *r*). NREM oscillation measures shown in the figures represent averages across electrodes in the significant cluster. Warm colors represent positive correlations and blue represents negative correlations. Electrodes marked in gray correspond to *p*_uncorrected_ < .05 and electrodes in magenta surpass cluster-level correction for multiple comparisons (*p*_corrected_ < .05). The right panels show scatterplots of these relationships in the NC (blue) and ANX (red) groups. The regression line is shown for the entire sample.

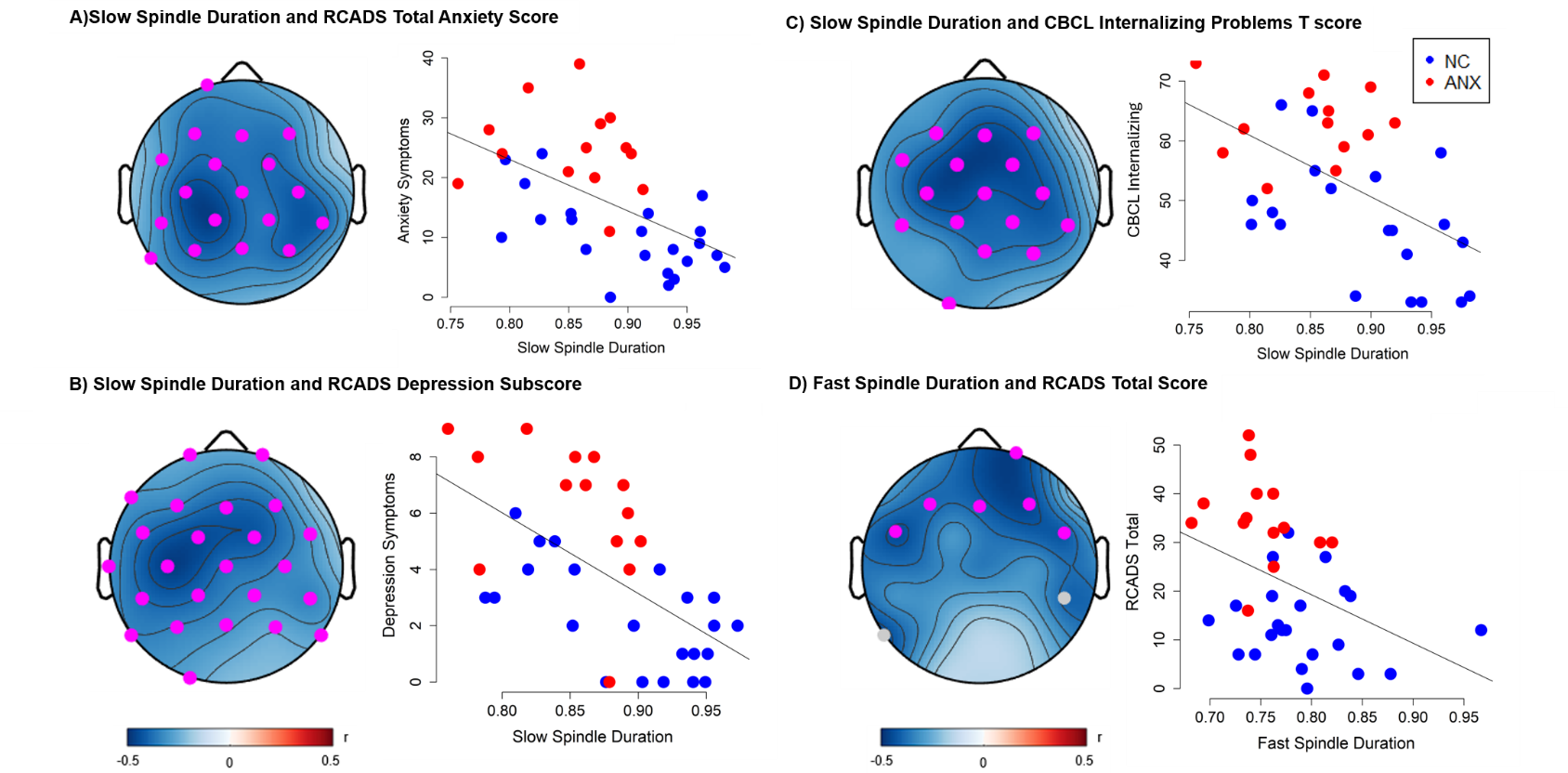

**Supplemental Figure 5. Associations of Slow Oscillation (SO) Density with Internalizing Symptoms.** SO density was positively associated with (A) RCADS Depression Symptoms, (B) RCADS Anxiety Symptoms, and (C) CBCL Internalizing Problems. Left panels show topographical maps of the correlation strength (Pearson's *r*). NREM oscillation measures shown in the figures represent averages across electrodes in the significant cluster. Warm colors represent positive correlations and blue represents negative correlations. Electrodes marked in gray correspond to *p*_uncorrected_ < .05 and electrodes in magenta surpass cluster-level correction for multiple comparisons (*p*_corrected_ < .05). The right panels show scatterplots of these relationships in the NC (blue) and ANX (red) groups. The regression line is shown for the entire sample.

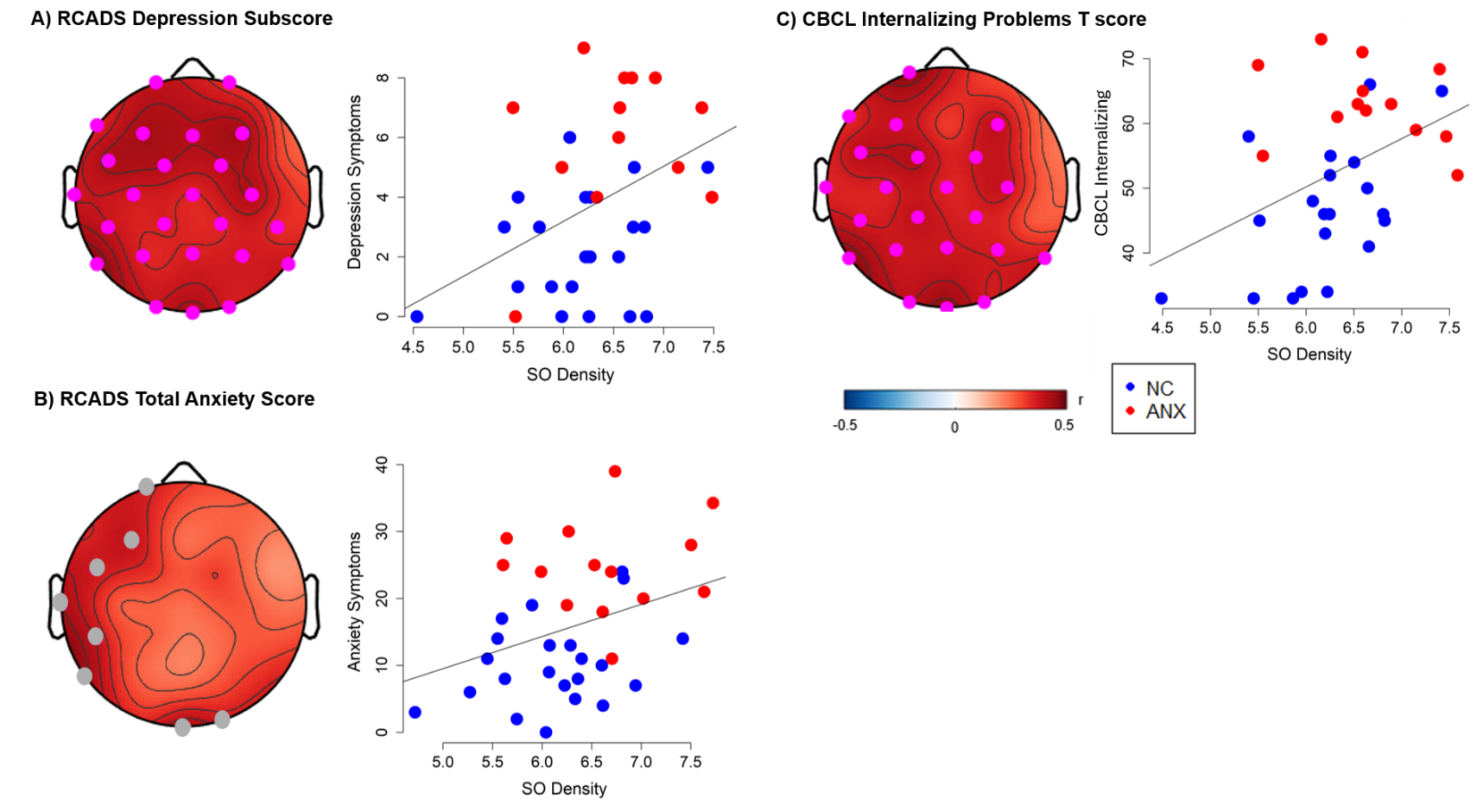

**Supplementary Table 1**. Patient diagnosis and medications. Participant numbers were changed to protect confidentiality. Information on medication use was unavailable for five ANX participants.

| Participant | Medication Class | Active Ingredient |
| --- | --- | --- |
| P1 | Antidepressant (SSRI) | Fluoxetine |
| P2 | Antidepressant (SSRI) | Escitalopram |
| P3 | Antidepressant (SSRI) | Sertraline |
| P4 | Not medicated |  |
| P5 | Antidepressant (SSRI) | Sertraline |
| P6 | Antidepressant (SSRI) | Escitalopram |
|  | Anxiolytic | Buspirone |
| P7 | Antidepressant (SSRI) | Sertraline |
| P8 | Antidepressant (SSRI) | Escitalopram |
| P9 | Antidepressant (SSRI) | Escitalopram |
| P10 | Antidepressant (SSRI) | Fluoxetine |
|  | Anxiolytic | Buspirone |
| P11 | Not medicated |  |
| P12 | Not medicated |  |

**Supplementary Table 2.** Linear Regression Analyses of NREM Oscillations with Cluster-Based Correction. The representative electrode corresponds to the electrode with the largest effect size within each significant cluster.

| Outcome | NREM Predictor | Cluster size | Corrected p | Representative Electrode | β | SE | f^2^ |
| --- | --- | --- | --- | --- | --- | --- | --- |
| RCADS Total | Slow Spindle Duration | 20 | .002 | C3 | -110.0 | 27.4 | .5 |
| RCADS Anxiety | Slow Spindle Duration | 18 | .002 | CP1 | -59.0 | 15.5 | .4 |
| RCADS Depression | Slow Spindle Duration | 24 | <.001 | C3 | -26.8 | 5.1 | .8 |
| CBCL Internalizing | Slow Spindle Duration | 15 | .01 | FC1 | -89.9 | 26.0 | .4 |
| RCADS Total | Fast Spindle Duration | 6 | .04 | Fp2 | -82.3 | 33.4 | .2 |
| RCADS Total | SO Density | 6 | .05 | P7 | 7.7 | 2.7 | .2 |
| RCADS Depression | SO Density | 25 | .01 | P7 | 1.8 | 0.6 | .3 |
| CBCL Internalizing | SO Density | 22 | .01 | P7 | 7.2 | 2.4 | .3 |
| Facial Arousal | Slow Spindle Duration | 7 | .04 | FC1 | -9.7 | 3.2 | .3 |
| Look Away # | Slow Spindle Duration | 9 | .03 | F3 | -4.6 | 1.8 | .2 |
